# RefDeduR: A text-normalization and decision-tree aided R package enabling accurate and high-throughput reference deduplication for large datasets

**DOI:** 10.1101/2022.09.29.510210

**Authors:** Jiaxian Shen, Fangqiong Ling, Erica M. Hartmann

## Abstract

As the scientific literature grows exponentially and research becomes increasingly interdisciplinary, accurate and high-throughput reference deduplication is vital in evidence synthesis studies (e.g., systematic reviews, meta-analyses) to ensure the completeness of datasets while reducing the manual screening burden. Existing tools fail to fulfill these emerging needs, as they are often labor-intensive, insufficient in accuracy, and limited to clinical fields. Here, we present RefDeduR, a text-normalization and decision-tree aided R package that enables accurate and high-throughput reference deduplication. We modularize the pipeline into text normalization, three-step exact matching, and two-step fuzzy matching processes. We also introduce a decision-tree algorithm, consider preprints when they co-exist with a peer-reviewed version, and provide actionable recommendations. Therefore, the tool is customizable, accurate, high-throughput, and practical. RefDeduR provides an effective solution to perform reference deduplication and represents a valuable advance in expanding the open-source toolkit to support evidence synthesis research.

## Introduction

As research becomes increasingly interdisciplinary, searching multiple platforms (e.g., PubMed, Web of Science) is vital to ensure the completeness of retrieved datasets in evidence synthesis studies, especially for systematic reviews and meta-analyses^1^. This makes reference deduplication a key step between search and screening^2^. Deduplication is usually accessible as a module in multiple tools along the evidence synthesis pipeline, including search platforms (e.g., Ovid), reference management software (e.g., EndNote, Zotero, synthesisr), and screening assistance tools (e.g., Covidence, Rayyan, Metta^2^, SRA-DM^3^, revtools^4^). Despite wide availability, existing modules are often labor-intensive, insufficient in accuracy, and limited to certain fields or databases (primarily clinical), probably because deduplication is only one element among their multiple functions. While still useful for small datasets, due to these limitations, many approaches quickly become impractical in the fast-growing era of big data. Already, systematic reviews take tens of weeks. It has been shown that a systematic review takes an average of 164 full-time equivalent days in environmental science^5^ and 67.3 weeks for a five-person team in medical fields^6^. Any processes that can accelerate computation and decrease manual labor are thus valuable improvements.

To address these challenges, we developed an R package, RefDeduR, specializing in reference deduplication. The pipeline is modularized into text normalization, three-step exact matching, and two-step fuzzy matching, making it highly customizable. With finely tuned text cleaning and normalization, RefDeduR’s high-confidence exact matching process outperforms many tools, even with their fuzzy matching procedure included. We semi-automate the time-consuming and error-prone manual review process in fuzzy matching by introducing a decision-tree algorithm, making it high-throughput while still maintaining accuracy. Additionally, we propose to use the inflection point of the similarity distribution curve as the cutoff threshold, making the pipeline more practical. The tool also takes into account preprints and conference proceedings, discarding them when a peer-reviewed version is present. This is and will be increasingly important with the rise of preprint servers. Last but not least, as a free open-source package, RefDeduR is highly interoperable with other software (including both commercial and open-source), will support the development of future tools (just like the various packages RefDeduR is built upon^4,7-10^), and more broadly will contribute to the prosperity of the evidence synthesis field.

Users can access RefDeduR on GitHub (https://github.com/jxshen311/RefDeduR) and view further documentation and examples on the website (https://jxshen311.github.io/RefDeduR/). Below, we demonstrate the functionality of RefDeduR with an example pipeline (see **Figure 1** for a flowchart of the recommended pipeline with key functions listed).

**FIGURE 1.**
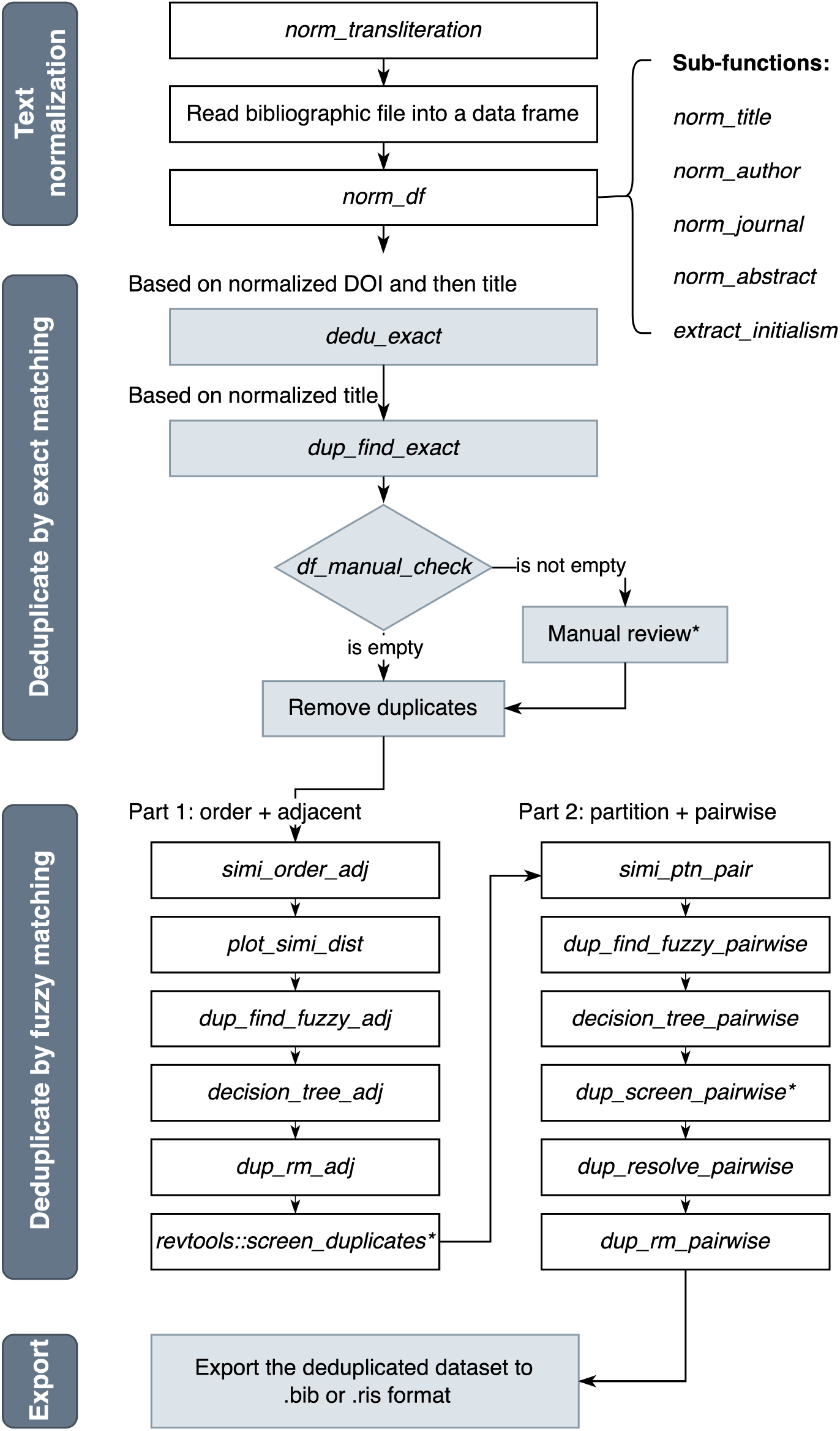
A flowchart of RefDeduR’s recommended pipeline with key functions listed. Functions are italicized to differentiate from the non-italicized descriptions. Processes involving manual review are marked with an asterisk. Refer to R documentation of each function and the tutorial (https://jxshen311.github.io/RefDeduR/articles/RefDeduR_tutorial.html) for more details.

## Functionality with an example pipeline

### Example dataset

The example dataset contains all bibliographic records (n = 6384) retrieved in a systematic review on indoor surface microbiome studies. We conducted the systematic search on 2022-01-10 through 3 platforms, and 1268, 3386, and 1733 records were retrieved from PubMed, Scopus, and Web of Science, respectively. The search terms consist of 4 key concepts: 1) indoor, 2) bacterial community/microbiome, 3) sequence-based, and 4) surface. Table 1 summarizes the number of records with missing values for relevant fields.

**TABLE 1.** Number of records with missing values for relevant fields

| Field | title | author | journal | abstract | year | DOI |
| --- | --- | --- | --- | --- | --- | --- |
| Number of records with missing values | 0 | 3 | 50 | 8 | 20 | 221 |

### Module 1: Text normalization

#### Transliterate non-ASCII characters

Before reading bibliographic files (usually BibTeX or RIS format) into data frames (the standard format for R datasets), we transliterate non-ASCII characters to the ASCII format using the function `norm_transliteration`. The default transliteration process includes: (1) transliterate common Greek letters to their names (e.g., “α” to “alpha”, “β” to “beta”) and (2) transliterate accented characters to ASCII equivalents (e.g., “á” to “a”, “ä” to “a”). Users can also perform transliteration based on customized rules or any other rules supported by `stringi::stri_trans_general`.

The transliteration increases the chance of successful deduplication by exact matching (**Module 2**). Additionally, it reduces noise when partitioning the dataset by the first two letters of `first_author_last_name_norm` at the fuzzy matching step (**Module 3**). For example, a record titled “Carriage and population genetics of extended spectrum **β**-lactamase-producing *Escherichia coli* in cats and dogs in New Zealand” sometimes has the title “Carriage and population genetics of extended spectrum **beta**-lactamase-producing *Escherichia coli* in cats and dogs in New Zealand”. Author names “Álvarez-Fraga, L. and Pérez, A.” are sometimes written as “Alvarez-Fraga, L. and Perez, A.”.

#### Read bibliographic file into a data frame

We leverage function `read_bibliography` from the R package revtools^4^ to read the transliterated bibliographic file (we recommend using BibTeX; see Github tutorial for reasons) into a data frame.

#### Text cleaning and normalization

Before deduplication, we perform multiple finely tuned text cleaning steps on the dataset. Text cleaning includes not only standard text normalization such as converting letters to lowercase, but also tailored operations in response to patterns we observed, such as removing trademark “(TM)” in title, removing English stop words in journal, and removing publisher/citation information in abstract (`norm_` functions, e.g., `norm_title`). Furthermore, we extract helper columns which we will use downstream (e.g., `journal_initialism` and `first_author_last_name_norm`). While the functions responsible for normalizing individual columns facilitate flexibility, a wrapper function, `norm_df`, is also provided to streamline the process. This function takes a data frame as the input and performs all normalization operations required in the downstream reference deduplication. Similarly, finer text normalization increases the probability of successful deduplication at the exact matching stage, where both accuracy and confidence are assured.

### Module 2: Deduplicate by exact matching

Following text normalization, we deduplicate by exact matching based on (1) normalized DOI (coded as `doi_norm`), (2) title, and (3) normalized title (`title_norm`) in order. DOI is chosen as the first metric because it is decisive (i.e., single selectivity). However, deduplication efficacy is limited by its medium/low applicability due to missing values. Hence, title is then employed, as it is high in both selectivity and applicability. Selectivity of a field equals 1 – (1/the number of unique field values). A field with high selectivity is one whose values are shared by only a limited number of records. A decisive field is a special case, where any non-null value is unique^2^. Applicability equals the number of non-null values divided by the number of total records, meaning that a high applicability field has few null values.

We use the function `dedu_exact` to automatically identify and remove duplicates. The user specifies one or multiple fields of the dataset according to which deduplication is conducted. If multiple fields are specified, deduplication will be performed one-by-one in order. Only records with non-null values will be investigated and the most recent version will be retained at removal. We recommend using `dedu_exact` for high-confidence fields such as normalized DOI and title.

Since completeness is crucial in evidence syntheses, we prioritize specificity over sensitivity for RefDeduR, and thus introduce a verification mechanism for fields that the user might consider less confident. Exemplified in the recommended pipeline, we use `dup_find_exact` to locate potential duplicates according to normalized title and check the outcome based on `first_author_last_name`. If the detected duplicate sets have different values in the verification field (in this case, `first_author_last_name`), they will be output for manual review. Typically, the number of duplicate sets requiring manual review is small at this step (e.g., in the example dataset, only 1 set needs to be reviewed). Note that incorporating the verification mechanism for normalized title is particularly conservative. If verification is not needed, the user can incorporate normalized title into `dedu_exact`. Moreover, although not included in the standard pipeline, the user can utilize the functions to further search for duplicates by exact matching other fields, such as normalized abstract.

### Module 3: Deduplicate by fuzzy matching

Once we remove all duplicates by the high-confidence exact matching processes, we proceed to fuzzy matching. Fuzzy matching is performed by calculating string similarity based on Levenshtein edit distance.

Two major practical challenges of making the fuzzy-matching process both accurate and high-throughput are (1) to choose a sensible cutoff threshold for the similarity score and (2) to reduce burden of manual review and accelerate the step. In RefDeduR, we propose two strategies to address these challenges. For challenge 1, we examine the similarity distribution plots and use the inflection point of the curve as the cutoff threshold. This value serves as a starting point to further finely-tune the threshold (discussed in more detail below). For challenge 2, we introduce a decision tree that incorporates multiple fields (e.g., title, author, year, journal, first author) to semi-automate the “manual review” step (illustrated in **Supplementary Figure 1**). This is especially helpful for large datasets, in which case the number of duplicate sets requiring manual review could be unfeasibly high (e.g., revtools outputs ∼1,400 duplicate sets for manual confirmation when treating this example dataset). Moreover, in addition to ensuring that the most recent record is retained at duplicate removal, we recognize the increasing prevalence of preprints and conference proceedings and remove them as well when co-existing with a peer-reviewed version.

To improve the computational efficiency, we divide this process into 2 parts: (1) order the records and compare only between the adjacent rows, and (2) perform pairwise comparisons between records within the same group after partitioning.

#### Part 1: order + adjacent

First, we calculate string similarity between adjacent rows for columns `title_norm` and `abstract_norm` using the function `simi_order_adj`. By default, the dataset is ordered alphabetically by `title_norm` before calculation, but the user may choose another field. Second, we plot similarity distributions of `title_norm` and `abstract_norm` by `plot_simi_dist` (Figure 2) to choose cutoffs. The plots suggest a cutoff score of 0.7 or 0.6 for the title and 0.3 for the abstract. For demonstration purpose, we use 0.7 and 0.3 here. The selected cutoffs are then passed to `dup_find_fuzzy_adj` to locate potential duplicates. The function outputs 2 data frames: (1) the input data frame with `match` column added and (2) a data frame listing `id` of duplicate pairs (`id_dup_pair_adj`). The decision tree is introduced next to semi-automate the “manual review” process. The function `decision_tree_adj` generates and adds decisions to `id_dup_pair_adj`. There are 3 possible levels of decisions: duplicate, not duplicate, and check. If the decision is not duplicate, the `match` column will be modified. To ensure a high accuracy, especially a low false positive rate, check is kept in the decision tree to signal manual confirmation. Finally, we deduplicate accordingly for different scenarios. For the duplicate, we remove duplicates directly by `dup_rm_adj`. For the check, we leverage `revtools::screen_duplicate`, leading to a graphical interface to interactively screen the duplicate pairs^4^.

**FIGURE 2.**
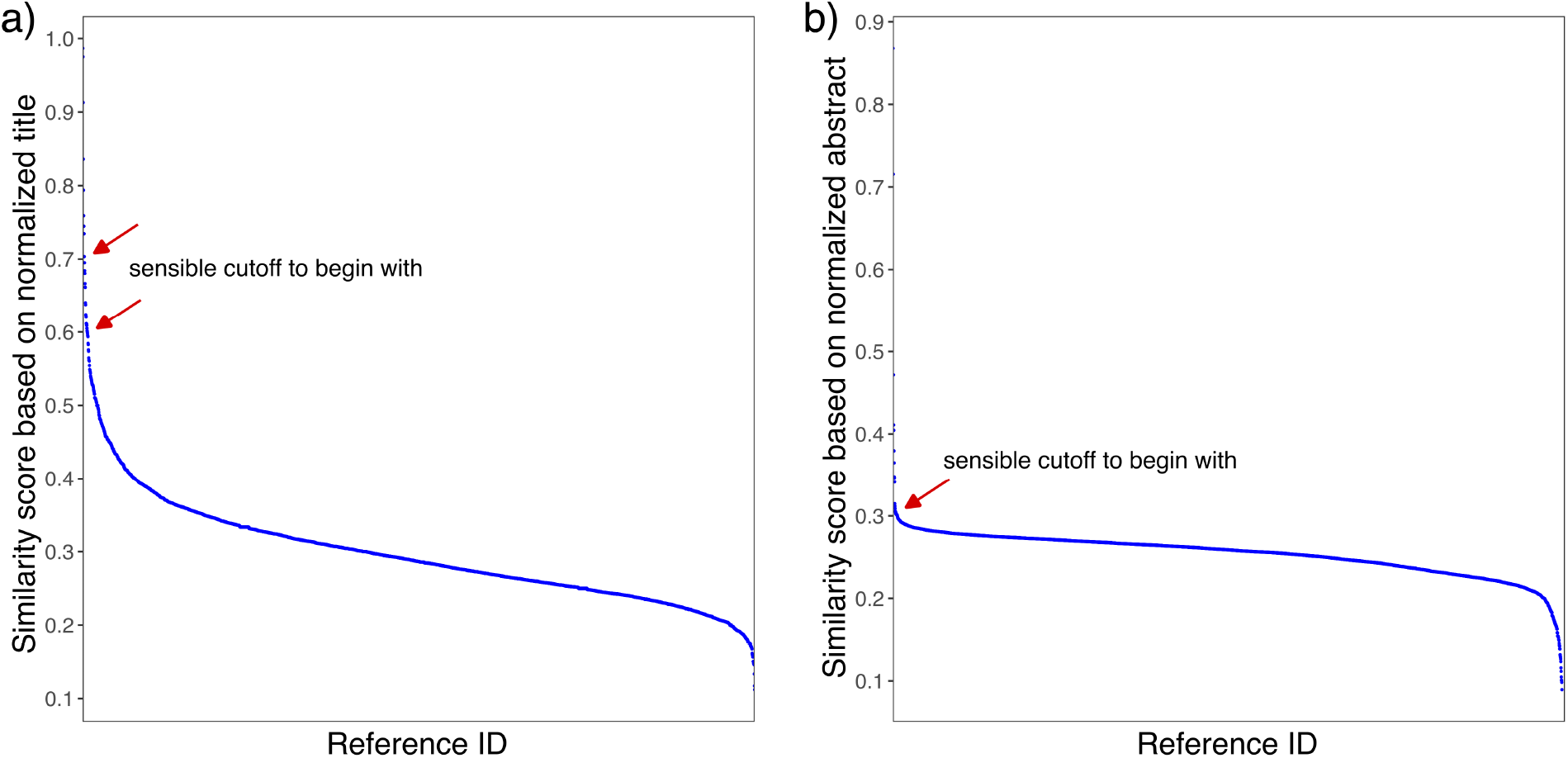
Distribution of string similarity scores of the example dataset based on a) normalized title and b) normalized abstract. Sensible cutoffs to begin with are marked by red arrows.

#### Part 2: partition + pairwise

We further deduplicate according to pairwise string similarity between all records within the same partitioned group. Following procedures similar to those in part 1, we first calculate string similarity for columns `title_norm` and `abstract_norm` using the function `simi_ptn_pair`. The difference is that we partition the dataset rather than order it. We recommend using the first two letters of `first_author_last_name_norm` as the partitioning metric, but the user has the flexibility to choose another parameter (e.g., year). We found the default metric more efficient than year for datasets that are skewed towards recent years. This is probably the case for many evidence synthesis studies, as the literature tends to grow exponentially. In addition, with the prevalence of preprints, partitioning by year becomes less accurate. Because the dataset is partitioned, results are now stored in lists as opposed to data frames in part 1. Likewise, we then locate potential duplicates by `dup_find_fuzzy_pairwise`. The cutoff thresholds can be inherited from part 1. To avoid over-deleting unique records, we suggest tightening the cutoff of abstract similarity to 0.7 (or 0.6) in this step, as opposed to 0.3 in part 1, where the risk is mitigated by the more restricted ordering. Decision tree is applied subsequently by `decision_tree_pairwise` and we can call `dup_screen_pairwise` to output the duplicate pairs that are labeled as check for manual review. Finally, we call `dup_resolve_pairwise` to resolve the check decisions to either duplicate or not duplicate accordingly and remove duplicates by `dup_rm_pairwise`.

We recommend using the inflection point as a data-driven threshold selection method based on the assumption that duplicates constitute a relatively small portion of the total after text normalization and exact matching. Thresholds from previous deduplication methods are largely empirical or anecdotal. For example, litsearchr uses “titles that are more than 95% similar, or abstracts that are more than 85% similar” but does not provide a theoretical basis for these cutoffs^11^. Compared with thresholds inherited from previous experience, this method is more quantitative. However, the selection of cutoff thresholds is based on visual observation of the distribution curves and does not actually need to be very precise. For example, the end result is not impacted by using 0.7 or 0.6 for the first threshold (**Supplementary Table 1**). Instead, numbers around that area all work similarly, partly due to the buffering functionality of the decision tree. Generally, a larger similarity score leads to higher specificity and may cause false negatives, while a smaller similarity score leads to higher sensitivity and may cause false positives. Regardless, all the 8 scenarios we tested had accuracy ≥ 99.94% (**Supplementary Table 1**; title similarity: 0.7 ∼ 0.5; abstract similarity: 0.7 ∼ 0.3). Specifically, the recommended schemes (S1 & S2) resulted in 0 false positive and 1 false negative. If the cutoff of abstract similarity was not raised to 0.7 (or 0.6) in part 2 (i.e., still used 0.3, S3 & S4), all duplicates would be identified at the cost of having 4 false positives. Towards the conservative direction, if 0.7 (or 0.6) was used throughout the procedures for both title and abstract similarity (S5 & S6), a false positive rate of 0 was maintained, and the number of false negatives was slightly increased to 3 for 0.7 and 2 for 0.6. Lowering the thresholds to 0.55 or 0.5 caused both false positives and false negatives (S7 & S8), but none exceeded 2.

### Export the deduplicated dataset

We can leverage `write_bibliography` from R package revtools^4^ to export the deduplicated data frame into a BibTeX or RIS file. Alternatively, R packages synthesisr (https://CRAN.R-project.org/package=synthesisr) and `RefManageR` (https://CRAN.R-project.org/package=RefManageR) also contain similar functions.

## Discussion

### Benchmarking

We benchmarked RefDeduR against existing tools using the example dataset. After manual curation, 3828 records were retained in the unique subset among the raw dataset with 6384 records (**Supplementary Table 2**). With this manually curated dataset as the benchmark set, deduplication performance was then assessed between March and September 2022. We considered deduplication modules from a variety of tools along the evidence synthesis pipeline, including search platforms (Ovid), reference management software (EndNote X20 v20.2, Zotero v6.0.9, Mendeley desktop v1.19.8 and synthesisr v0.3.0), and screening assistance tools (Covidence, Rayyan, Metta^2^, SRA-DM^3^, and revtools v0.4.1^4^). Ovid, Metta, and SRA-DM were excluded after a preliminary examination because their functionality was restricted to clinical databases. Deduplication of the dataset was then performed for the other software. We used default settings for Endnote X20, Covidence, Zotero, Mendeley and Rayyan, and chose the highest-performance scenario for revtools and synthesisr since they provided multiple options in their documentation. Version information is not available for Covidence and Rayyan but the operation date as well as other details are described at https://github.com/jxshen311/RefDeduR_benchmark.

Quantitative evaluation was not obtained using Mendeley or Rayyan due to the lack of an option to automatically resolve detected duplicates. Mendeley desktop found 1457 sets of duplicates. Rayyan detected 3369 potential duplicates out of the raw dataset. Both are unfeasibly labor intensive with manual resolution as the only option. Moreover, according to the official announcement, Mendeley desktop has been discontinued and will be gradually replaced by the web-centric version, Mendeley Reference Manager, which does not currently support reference deduplication. Mendeley desktop also had a worse performance than Covidence in a previous comparison study based on a smaller dataset (n = 3130), which can be used as an indirect indicator^12^. While Rayyan claims that it automatically detects and resolves 100% duplicate articles, none of the flagged duplicates were automatically deleted in the example dataset. Despite failure to make a quantitative comparison, we uploaded the dataset that had gone through RefDeduR’s deduplication for Rayyan to further perform deduplication. It found 8 duplicates, all of which were deemed false positives by manual review. This indicates that Rayyan has no higher sensitivity and lower specificity than RefDeduR.

The remaining tools were evaluated quantitatively. We first identified false positives and false negatives, and then calculated accuracy, sensitivity, and specificity accordingly as described previously (**Supplementary Table 3**)^12^. All these tools except Zotero support automatic resolution, for which user-developed scripts are available as interim workarounds. For example, a developer with the user name “marcelparciak” posted a Java script that automates the clicking of “Merge X items” button for 100 times with a second waiting time in between on the Zotero forum (https://forums.zotero.org/discussion/40457/merge-all-duplicates). Although in practice, the script stopped frequently (after ∼ 10 clicks) and required a manual restart, we resolved all duplicates after approximately a day.

RefDeduR deduplicated the dataset to 3829 records, with only one record missed in comparison to the benchmark set, while all the other tools missed substantially more duplicate records for exclusion (i.e., false negatives) (**Figure 3**). Notably, with only the exact matching module, RefDeduR has already outperformed all the benchmarked tools, possibly due to the optimized text normalization. All the tools had 100% specificity for the example dataset except revtools (99.97%), which mis-identified one record as duplicate (i.e., false positive). This is surprising for Endnote and Zotero since their specificity was previously reported to be only 89% and 99%, respectively^12^. This means 208 and 20 false positives out of a dataset with 3130 references. It is possible that the software has enhanced their deduplication capability since this previous study that used Endnote X9 and was conducted between December 2018 and January 2020. Alternatively, variation between the two benchmark sets may also contribute to the different outcomes. In contrast to searching 3 platforms (PubMed, Scopus, and Web of Science) in our study, the benchmark set in the previous study was retrieved from Ovid only and was mostly clinically focused.

**FIGURE 3.**
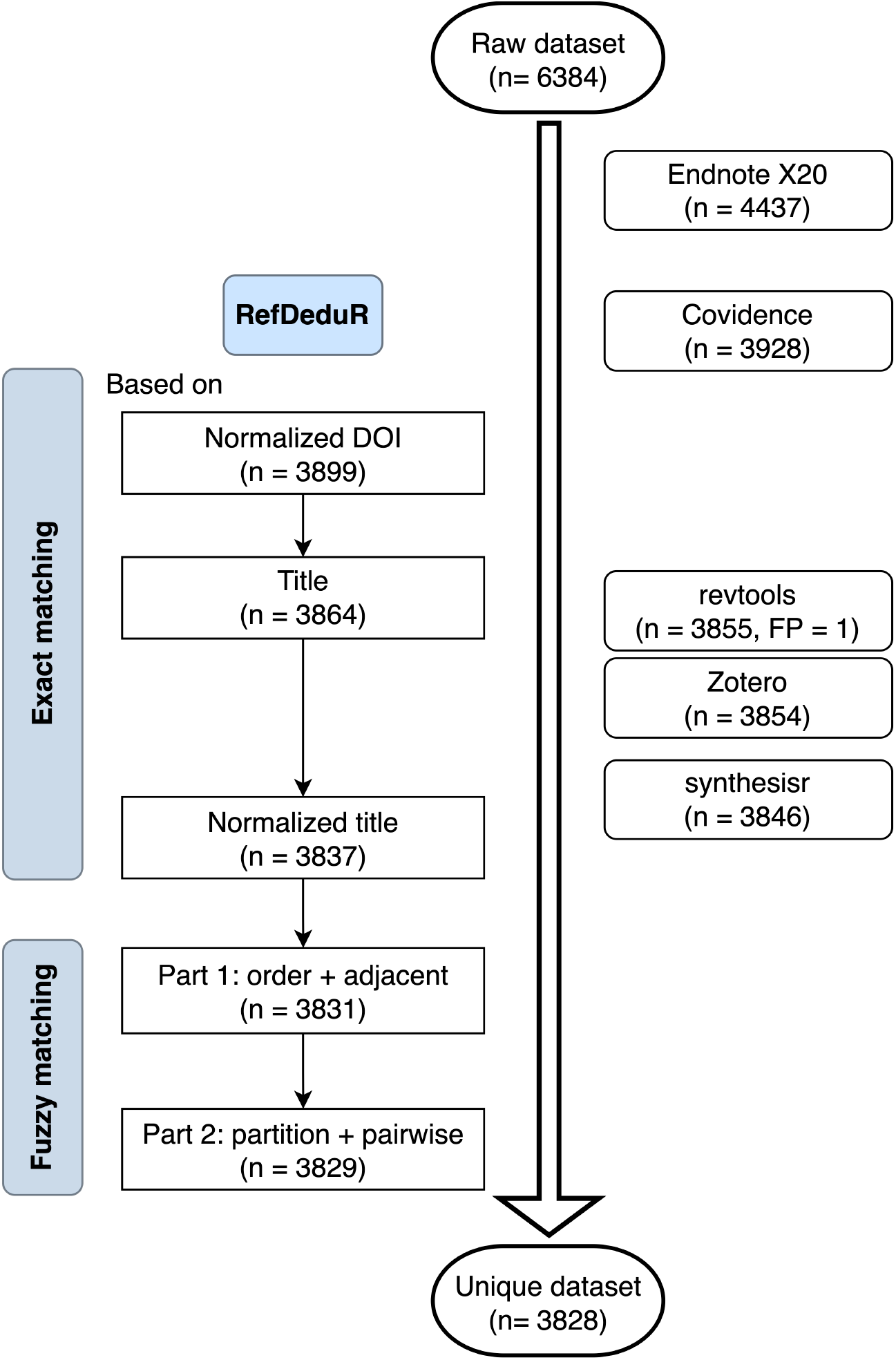
Comparison of deduplication performance between RefDeduR and existing tools. Performance of intermediate compartments is also displayed for RefDeduR. Default settings were used for Endnote X20, Covidence, and Zotero. The tools offering multiple options (e.g., revtools and synthesisr) are represented by the highest-performance one. Except revtools, all the other tools have 0 false positives (FP) (i.e., 100% specificity) for the example dataset.

RefDeduR balances risk of misidentification and manual burden via modularizing the entire process and combining the automatic decision-tree algorithm with the attenuated manual review. In contrast, the other tools rely heavily on users to check the flagged duplicates by design, thus limiting their robustness. Following the recommended pipeline of RefDeduR, only 3 duplicate sets needed to be reviewed before achieving the manually assured result (1 in the exact matching step and 2 in part 1 of the fuzzy matching step). However, the user had to screen 1460 duplicate sets for revtools and 556 for synthesisr. The numbers of duplicate sets requiring manual review were not disclosed by Endnote X20, Covidence, and Zotero. Nevertheless, they were approximated to be 974, 1228, and 1265 assuming all were duplicate pairs. In addition, because of the decision tree, RefDeduR’s manual workload is less susceptible to the change of fuzzy-matching thresholds. For instance, after lowering the similarity threshold from 0.7 to 0.6, 22 more duplicate sets were inputted into the decision tree (from 35 to 57), while the manual workload only increased by 6 (from 2 to 8) (**Supplementary Table 1**).

### General recommendations, potential limitations, and future directions

We recommend following the example pipeline when using RefDeduR, but users are offered the flexibility to build a custom pipeline. For example, if the user is satisfied with the number of output records, they may stop after the exact-matching module or part 1 of the fuzzy-matching module, since these modules are substantially faster (< 1 min) than part 2 of the fuzzy-matching module (∼20 min for an exhaustive similarity calculation). Alternatively, combining the RefDeduR’s intermediate or final output with other tools (e.g., Rayyan, revtools) could be a reassuring operation to further increase the possibility of achieving a desired outcome. In this way, the heavy manual review burden of these tools could also be relieved. For instance, using RefDeduR’s intermediate output from part 1 of the fuzzy-matching module as the input for Rayyan led to only 10 duplicates for manual review, as opposed to 3369 if the raw dataset was inputted.

Future versions of RefDeduR will further expand automated matching to focus on accuracy without increasing the burden on the user. For example, the decision tree could be expanded to include more fields (e.g., volume, issue and page). Due to the difficulty of obtaining a manually confirmed unique dataset, RefDeduR was only benchmarked using one dataset at this stage. We will continue improving the tool as more data become available. For instance, it is interesting to further explore the impact of threshold selection on software performance and the efficacy of training machine learning models for duplicate classification.

In conclusion, RefDeduR provides an effective solution to perform reference deduplication and represents a valuable advance in expanding the open-source toolkit to support evidence synthesis research. It will also support the development of future tools, just like the packages RefDeduR is built upon (e.g., revtools).

## Data availability

Users can access RefDeduR on GitHub (https://github.com/jxshen311/RefDeduR) and view further documentation and examples on the website (https://jxshen311.github.io/RefDeduR/). A step-by-step tutorial is available at https://jxshen311.github.io/RefDeduR/articles/RefDeduR_tutorial.html. Source code, supplementary data, and additional descriptions about the benchmarking analysis are available at https://github.com/jxshen311/RefDeduR_benchmark.

## Funding

This research was supported by a Terminal Year Fellowship from the Richter Memorial Fund to JS.

## Conflict of Interest

The authors declare that the research was conducted in the absence of any commercial or financial relationships that could be construed as a potential conflict of interest.

## Acknowledgments

We thank Yutong Wu for the illuminating discussions about the design of RefDeduR. We are also grateful to Ruochen Jiao and Alexander G. McFarland for their help in coding.

## Highlights

- Accurate and high-throughput reference deduplication is critical in evidence synthesis studies (e.g., systematic reviews).
- While this functionality is available in many tools, they are gradually outdated in the fast-growing era of big-data and interdisciplinary research, since they are often labor-intensive, insufficient in accuracy, and limited to clinical fields.
- RefDeduR is a text-normalization and decision-tree aided R package that enables accurate and high-throughput reference deduplication.
- The software is highly customizable and practical with actionable recommendations provided. Additionally, it takes preprints into account in deduplication.
- The tool enhances the power of reference deduplication in response to emerging needs and is especially helpful for large datasets.

## Notes

https://github.com/jxshen311/RefDeduR

https://jxshen311.github.io/RefDeduR/

https://github.com/jxshen311/RefDeduR_benchmark

### Competing Interest Statement

The authors have declared no competing interest.

