## Supplementary Figure 1 for "RefDeduR: A text-normalization and decision-tree aided R package enabling accurate and high-throughput reference deduplication for large datasets"

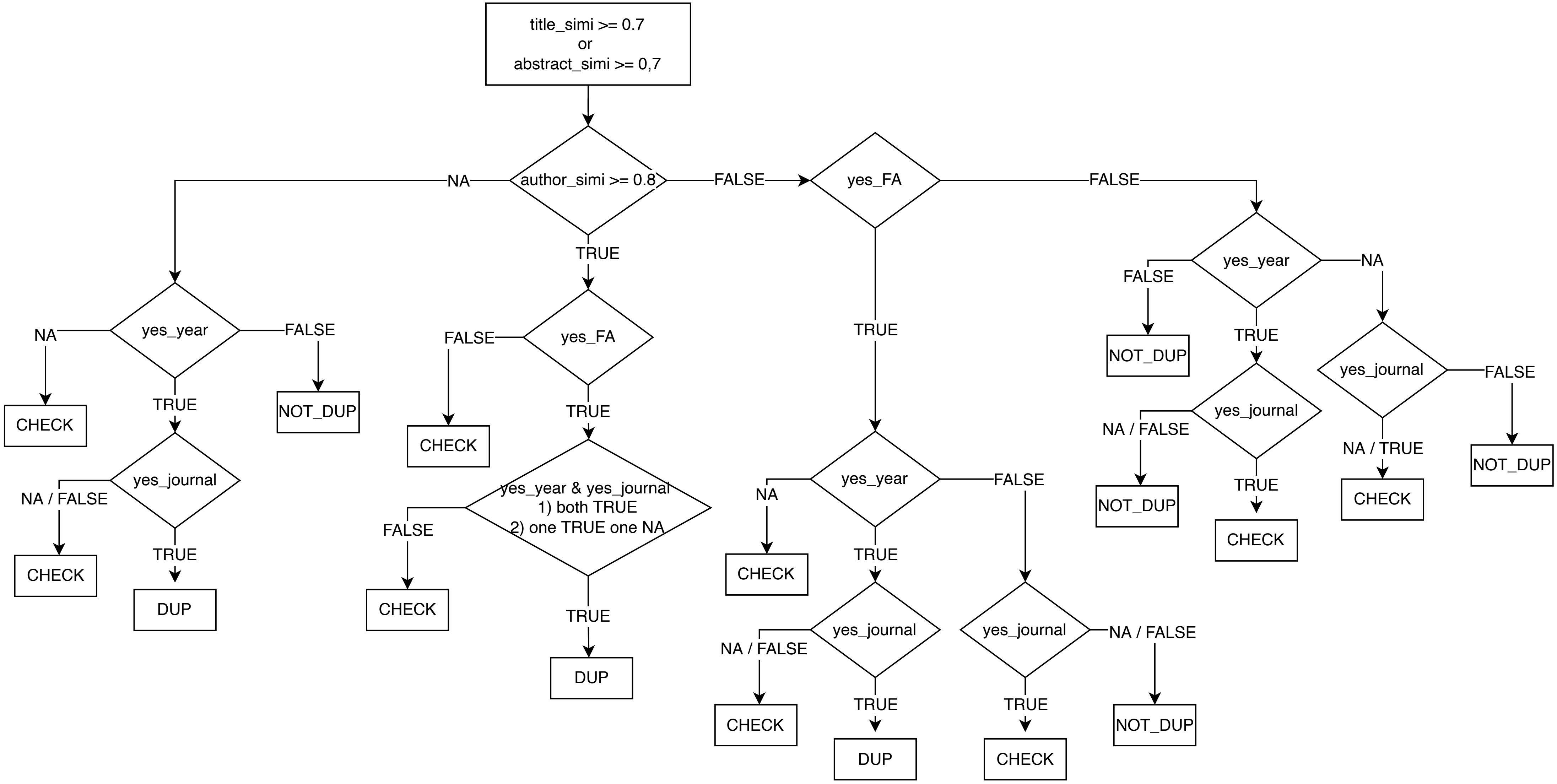


**Supplementary Figure 1** A detailed illustration of the decision tree used in fuzzy-matching processes.

**List of abbreviations:**

| Abbreviation | Explanation | Abbreviation | Explanation |
|---|---|---|---|
| title_simi | title similarity | yes_journal | journal initialism is identical or one of the records is a preprint or conference proceeding |
| abstract_simi | abstract similarity | | |
| author_simi | author similarity | CHECK | decision is "check" |
| yes_FA | normalized first_author_last_name is identical | DUP | decision is "duplicate" |
| yes_year | year is identical or one of the records is a preprint or conference proceeding | NOT_DUP | decision is "not duplicate" |
