## Supplementary Table 1 for "RefDeduR: A text-normalization and decision-tree aided R package enabling accurate and high-throughput reference deduplication for large datasets"

**Supplementary Table 1** Comparison of RefDeduR's performance for different similarity thresholds.

| | | Recommended | | Risky<br>Cause false positives | | Conservative<br>Slightly more false negatives | | Less accurate<br>One or two misidentifications | |
|---|---|---|---|---|---|---|---|---|---|
| **Scenario** | | **S1** | **S2** | **S3** | **S4** | **S5** | **S6** | **S7** | **S8** |
| Number of records after exact matching | | 3837 | 3837 | 3837 | 3837 | 3837 | 3837 | 3837 | 3837 |
| Fuzzy matching part 1: threshold for title similarity | | 0.7 | 0.6 | 0.7 | 0.6 | 0.7 | 0.6 | 0.5 | 0.55 |
| Fuzzy matching part 1: threshold for abstract similarity | | 0.3 | 0.3 | 0.3 | 0.3 | 0.7 | 0.6 | 0.5 | 0.55 |
| Number of duplicate pairs entering the decision tree | | 30 | 41 | 30 | 41 | 9 | 26 | 86 | 39 |
| Automatic decision | check | 2 | 4 | 2 | 4 | 1 | 3 | 4 | 3 |
| | duplicate | 5 | 5 | 5 | 5 | 3 | 4 | 4 | 4 |
| | not duplicate | 23 | 32 | 23 | 32 | 5 | 19 | 78 | 32 |
| Manual decision for "check" | duplicate | 1 | 1 | 1 | 1 | 1 | 1 | 1 | 1 |
| | not duplicate | 1 | 3 | 1 | 3 | 0 | 2 | 3 | 2 |
| Number of records after part 1 | | 3831 | 3831 | 3831 | 3831 | 3833 | 3832 | 3832 | 3832 |
| Fuzzy matching part 2: threshold for title similarity | | 0.7 | 0.6 | 0.7 | 0.6 | 0.7 | 0.6 | 0.5 | 0.55 |
| Fuzzy matching part 2: threshold for abstract similarity | | 0.7 | 0.6 | 0.3 | 0.3 | 0.7 | 0.6 | 0.5 | 0.55 |
| Number of duplicate pairs entering the decision tree | | 5 | 16 | 92 | 98 | 5 | 16 | 42 | 28 |
| Automatic decision | check | 0 | 4 | 24 | 25 | 0 | 4 | 9 | 6 |
| | duplicate | 2 | 2 | 7 | 7 | 2 | 2 | 4 | 3 |
| | not duplicate | 3 | 10 | 61 | 66 | 3 | 10 | 29 | 19 |
| Manual decision for "check" | duplicate | 0 | 0 | 0 | 0 | 0 | 0 | 0 | 0 |
| | not duplicate | 0 | 4 | 24 | 25 | 0 | 4 | 9 | 6 |
| Number of records after part 2 (Final outcome) | | 3829 | 3829 | 3824 | 3824 | 3831 | 3830 | 3828 | 3829 |
| Performance statistic | False positive | 0 | 0 | 4 | 4 | 0 | 0 | 2 | 1 |
| | False negative | 1 | 1 | 0 | 0 | 3 | 2 | 2 | 2 |
| | Accuracy (%) | 99.98 | 99.98 | 99.94 | 99.94 | 99.95 | 99.97 | 99.94 | 99.95 |
| | Specificity (%) | 100 | 100 | 99.9 | 99.9 | 100 | 100 | 99.95 | 99.97 |
| | Sensitivity (%) | 99.96 | 99.96 | 100 | 100 | 99.88 | 99.92 | 99.92 | 99.92 |
